# Regulation of vocal precision by local noradrenergic modulation of a motor nucleus

**DOI:** 10.1101/218479

**Authors:** Christopher M Glaze, Christina B Castelino, Steve P Bibu, Elvina Yau, Marc F Schmidt

**Affiliations:** Biology Department, University of Pennsylvania, Philadelphia, PA 19104; Neuroscience Graduate Group, University of Pennsylvania, Philadelphia, PA 19104; Biological Basis of Behavior Program, University of Pennsylvania, Philadelphia, PA 19104; <u>Current Address</u>: The Lockwood Group, Stamford, CT 06901.; <u>Current Address</u>: Department of Neuroscience, University of Pennsylvania, Philadelphia, PA 19104

**Author notes:** <u>Corresponding Author</u>: <u>Email</u>. Both authors contributed equally to this work.

## Abstract

Recent theories of norepinephrine (NE) function suggest that NE modulates the transition between stereotyped, goal-directed behavior and more variable exploratory behaviors that facilitate learning and adaptation. We provide evidence for context dependent switching by NE that is analogous to this explore/exploit strategy in the vocal system of the zebra finch (*Taeniopygia guttata*). Stimulation of the locus coeruleus, the major source of norepinephrine in the brain, decreases song trial-to-trial variability transforming the variable, exploratory “undirected” song into song that resembles the more stereotyped, exploitative “directed” song males sing to females. This behavioral switch is mediated by NE acting directly on a cortical motor nucleus that integrates inputs from a premotor cortical nucleus and a basal ganglia circuit necessary for vocal motor learning. These findings suggest that norepinephrine can act directly on the motor system to influence the transition between exploratory and exploitative behavioral strategies.

## INTRODUCTION

Activation of the locus coeruleus (LC), through its diffuse projections, has been traditionally implicated in regulating arousal levels (Kety, 1972) and modulating neural responsiveness in sensory and learning circuits (Hasselmo et al., 1997; Berridge and Waterhouse, 2003). More recent theoretical work has suggested that LC mediated release of norepinephrine (NE) may facilitate the optimization of behavior with respect to rewards and goals by increasing the gain of neural activation functions, directly affecting the precision of action selection among a discrete set of options in learning and decision-making tasks (Aston-Jones and Cohen, 2005a). According to this proposal, the noradrenergic system modulates behavioral flexibility, controlling the switch between the exploration of many different actions and the exploitation of those specific actions that yield the greatest utility (Aston-Jones and Cohen, 2005a, b). This theory requires that noradrenergic projections have a great degree of target specificity. Support for this idea comes from recent findings suggesting that LC, rather than containing a homogeneous population of neurons as previously thought, is compartmentalized into distinct computational units (Chandler et al., 2014b) that each might have the capability to independently modulate select cortical areas (Mather et al., 2015; Chandler, 2016). In the present study we test the hypothesis that norepinephrine plays a causal role in action selection in a continuous motor space and that it exerts its effects by targeting a single motor cortical nucleus in the songbird vocal control circuit.

Birdsong consists of precisely produced sequences of motor gestures that contain complex acoustic spectrotemporal structure on multiple timescales (Glaze and Troyer, 2006; Amador et al., 2013). Male zebra finches learn their song by adulthood, but even as adults they continue to alternate between highly precise “directed” song performed in the presence of a female and more variable, possibly exploratory, “undirected” song that is produced when singing alone (Sossinka and Boehner, 1980; Kao et al., 2005). Directed singing can be thought of as an exploitative behavior, where song is a precisely produced motor performance directed at the female (Tumer and Brainard, 2007; Andalman and Fee, 2009; Sober and Brainard, 2009; Ali et al., 2013). The more variable, undirected song has been suggested to promote active exploration of a motor space that facilitates learning of potentially new acoustic features (Tumer and Brainard, 2007; Sober and Brainard, 2009; Olveczky et al., 2011). The switch between the directed and undirected song types is mediated by a specialized basal ganglia circuit, known as the anterior forebrain pathway (AFP) (Figure 2a) (Farries, 2004; Reiner et al., 2004; Kao and Brainard, 2006; Farries and Perkel, 2008; Fee and Goldberg, 2011). Experimental inactivation of the AFP output nucleus LMAN causes the normally variable undirected song to become highly stereotyped and therefore resemble the acoustically precise directed song (Kao et al., 2005).

The locus coeruleus is highly conserved across vertebrates (Smeets and Gonzalez, 2000) and, in songbirds, it sends afferents throughout the “song control circuit” (Castelino and Schmidt, 2010). While NE transmission has been shown to modulate auditory response properties (Cardin and Schmidt, 2004; Castelino and Schmidt, 2010), its role in vocal production remains undefined (Hara et al., 2007). A key recipient of locus coeruleus input is the motor “cortical” area RA, a structure that is necessary for song production and serves as the major target of LMAN, the primary output nucleus of the AFP (Castelino and Schmidt, 2010). In vitro slice work provides indirect evidence that NE may influence vocal production because it can selectively suppress LMAN inputs onto RA without changing the strength of synaptic inputs from HVC, a premotor nucleus necessary for song production (Sizemore and Perkel, 2008). Noradrenergic release from LC terminals in RA therefore is well positioned to gate inputs from a basal ganglia circuit that is implicated in the generation of acoustic variability during undirected song (Kao et al., 2005; Kao and Brainard, 2006; Andalman and Fee, 2009).

Here we show evidence for direct influence of the noradrenergic system on acoustic variability through a combination of direct electrical stimulation of LC and reversible infusion of NE into RA in awake, freely moving birds.

## RESULTS

### Quantification of differences between directed and undirected song variability and singing rate

To capture the full extent that social context, or manipulation of the noradrenergic system, might have on the spectro-temporal properties of song in the zebra finch, we developed a novel “Euclidean variability” measure that uses average Euclidean distance between syllables and respective time-warped templates in the spectral domain. Prior studies investigating changes in song variability between directed and undirected song relied primarily on changes in variability in the fundamental frequency of individual syllable portions (Kao and Brainard, 2006; Andalman and Fee, 2009), which limits analysis to a small subset of song elements that happen to be measurable in that feature space (i.e. have a clear fundamental frequency to begin with). Our more general measure was not restricted to any particular kind of song element, and we verified that our novel methodological approach both captured variance in fundamental frequency (figure 1c) and, as shown below, could replicate previous results by analyzing song in adult male zebra finches both under conditions of undirected song (male singing alone) and directed song (male singing in the presence of a female).

**Figure 1.**
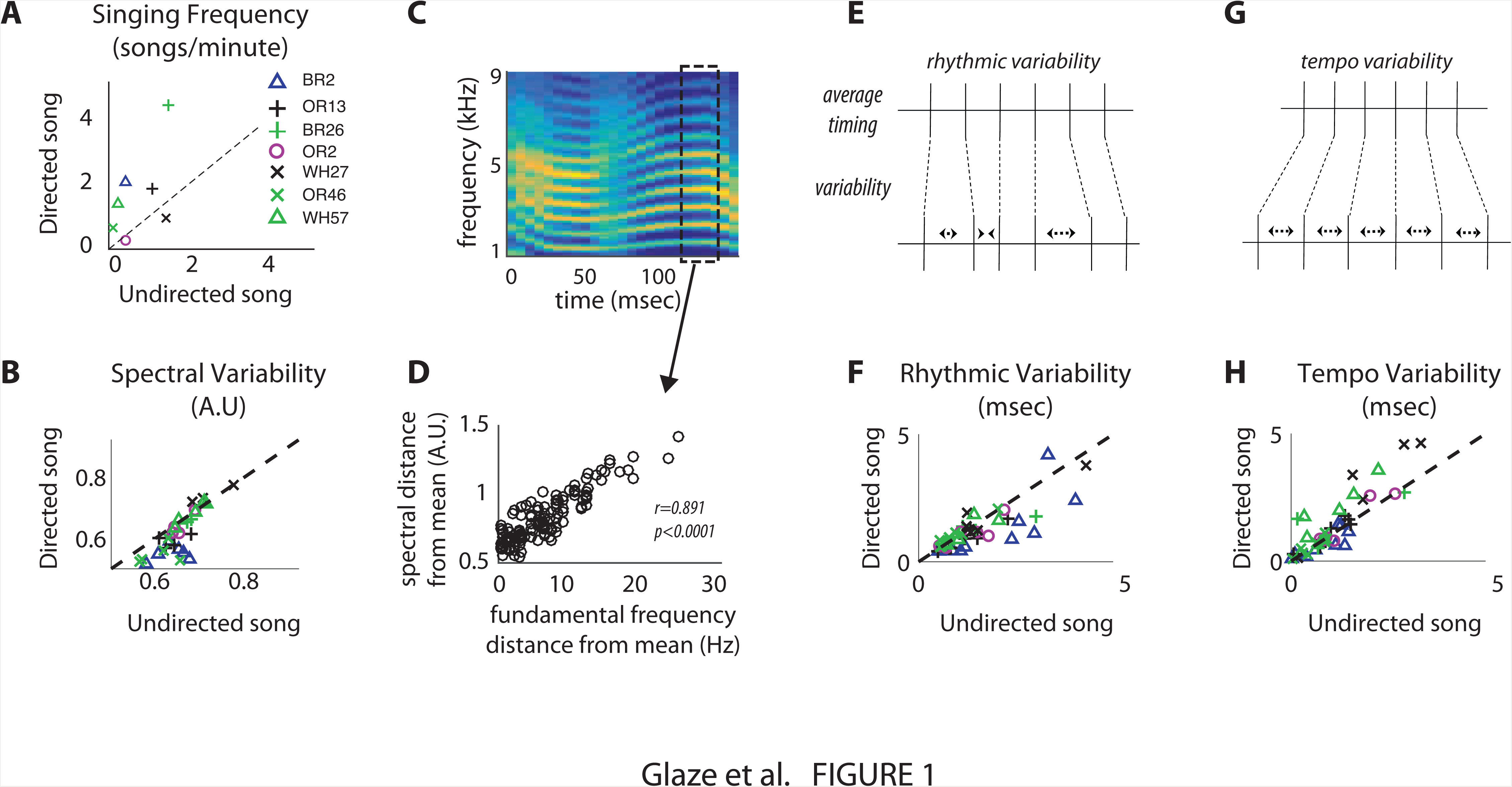
**Context-dependence of the behavioral parameters extracted for analysis.** (A) Scatterplot of the average number of songs produced per minute of recording from each bird by context. Measures taken from the female-present context (directed song) fall along the vertical axis and those taken from alone-context (undirected song) fall along the horizontal axis. Each bird is indicated by different symbols or colors. (B) Scatterplot of the total amount of trial-to-trial spectral variability in each song syllable by context (34 syllables in 7 birds). Vertical axis is the female-present context and the horizontal axis male-alone context. Units are arbitrary and therefore indicated as A.U. (Arbitrary Units). Syllables from individual birds are represented by different symbols or colors (C, D): Validation of the novel analytical method used to compute the variability measures in (B) (see Methods for details). (C) Segment of a song syllable from one bird with a stable and clearly defined fundamental frequency, rendering it amenable to the standard analysis of variability as in previous investigations into context-dependence of trial-to-trial spectral variability. (D) Scatterplot of spectral distance measured using the novel method on the vertical-axis vs. fundamental frequency deviations from the average along the horizontal axis for the same syllable segment indicated in (C). Data are plotted for 498 renditions of the same syllable. (E-H) Trial-to-trial timing variability model and context-dependent parameter estimates. (E) Schematic illustrating an example of rhythmic variability, in which trial-to-trial deviations in the time intervals between identified song features (e.g. syllable onsets and offsets) are independent but significantly affect overall song duration. (F) Scatterplot of the total amount of trial-to-trial rhythmic variability in each song syllable by context. (G) Schematic illustrating an example of tempo variability, in which trial-to-trial deviations in the time intervals between identified song features are correlated, preserving the relative durations of the time intervals much more than rhythmic variability. (H) Scatterplot of the total amount of trial-to-trial tempo variability in each song syllable by context, with axes as in (A, B, F). All birds used in these analyses were the same as those used for the infusion experiment (Figures 4, 5 and 6).

Songs were recorded from seven adult male zebra finches under both directed and undirected conditions, with motifs typically consisting of 3 – 9 syllables (median=5). Using our more general spectral variability measurement, we confirmed that syllables produced during directed singing have significantly less spectral variability than those produced during undirected singing, with decreases in variability estimates for each syllable of 5.64 ± 2.06% (figure 1b; hierarchical bootstrap t-test, p<0.005) with that direction of change holding in 6 of 7 birds. Variability differences appeared to hold across a wide variety of song syllables, many with complex structure that would not have been measurable with previous feature-based methods, indicating that context-dependent differences are generally spread across different song elements and not restricted to harmonic stack syllables with well-defined fundamental frequencies.

From this same dataset we also measured changes in song rhythmic (figure 1e) and tempo (figure 1g) variability during directed and undirected song using a statistical timing variability model previously shown to make principled definitions and estimates of these measures (Glaze & Troyer, 2013, 2010) . Specifically, rhythmic variability consists of rendition-to-rendition timing changes that are uncorrelated across a sequence of syllables and silent gaps between syllables in a motif, but nonetheless influence overall motif duration (unlike measurement error). In contrast, tempo variability measures the way in which all song elements in a given rendition become stretched or compressed in tandem. Interestingly, we found significant decreases in rhythmic variability (figure 1f) in the presence of a female with changes in this measure on a syllable-by-syllable and gap-by-gap basis of 12.48±7.66% (hierarchical bootstrap t-test, p<0.005), with the direction of change holding in 6 of 7 birds. By contrast, tempo variability showed an increase of 56.44±26.00% (hierarchical bootstrap t-test, p<0.05; figure 1h), with that direction of change persisting in 5 of 7 birds. As shown in previous studies (Sossinka and Boehner, 1980; Glaze and Troyer, 2006), there was a trend towards duration decreasing (tempo increasing) across time intervals in the presence of a female, with an average change of 0.4±0.4% that was consistent in 5 of 7 birds although it failed to reach significance (hierarchical bootstrap t-test, *p*=0.13).

Finally, we also corroborated previously reported findings on the influence of female presence on song rate, with the presence of a female increasing the rate of vocal sequence production by an average of 1.372±0.63 songs/minute (hierarchical bootstrap t-test, p=0.009), with the direction of change holding in 5 of 7 birds (figure 1a).

### Locus Coeruleus stimulation increases song rate and causes a decrease in song spectral and rhythmic variability

To examine the influence of the noradrenergic system on song production and variability parameters, we surgically implanted stimulating wires in LC (figure 2b) and quantified song features in the presence and absence of LC stimulation (N=4 birds). Specifically, over a period of many days (3 – 12 days), we compared songs produced under non-stimulated control conditions with songs produced during 1-Hz tonic stimulation of LC. Because we specifically were interested in whether an increase in noradrenergic tone caused by stimulation of LC would decrease the variability observed during undirected song, all of the recordings in this first set of experiments were performed on males singing undirected song in the absence of females. Experiments consisted of periods of LC stimulation (1-2 hours) interleaved with periods of non-stimulation of roughly the same duration. Typical daily sessions consisted of 4-6 interleaved blocks, the order of which was alternated every other day (Figure 3a).

**Figure 2.**
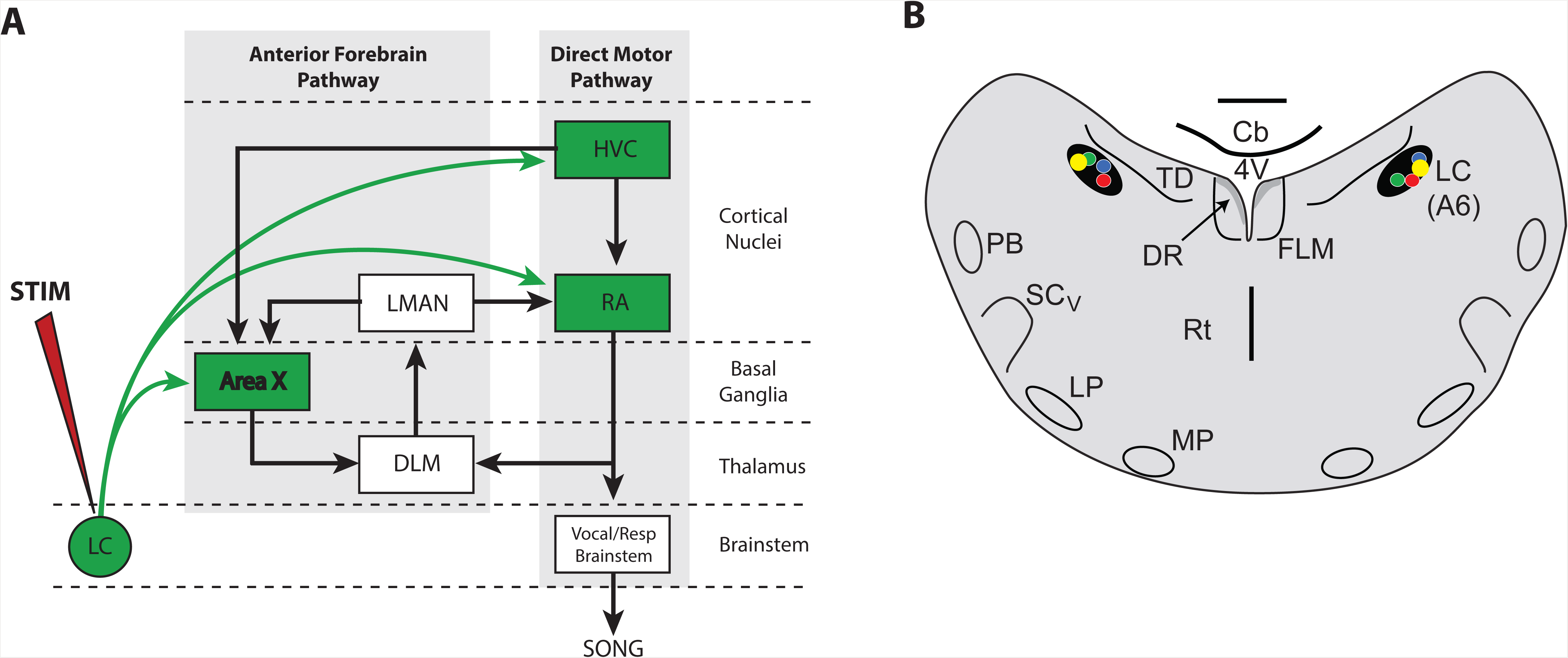
**Locus Coeruleus projections to the song control system.** We used stimulating electrodes placed bilaterally in the locus coeruleus (LC) to test the effect of tonic 1Hz stimulation on spectral variability of undirected song. (A) Schematic illustration of the known projections of LC onto HVC (Appeltants et al., 2001), RA (Appeltants et al., 2002), and area X (Castelino et al., 2007) in the song control system. (B) Coronal section of the midbrain showing successful bilateral placement of stimulating electrodes in LC. Individual electrode placement is illustrated by blue, green, yellow and red dots. <u>Anatomical abbreviations</u>: LMAN: lateral magnocellular nucleus of the anterior nidopallium; DLM: medial nucleus of the dorsolateral thalamus; RA: robust nucleus of the arcopallium ; HVC: used as proper name; Cb: cerebellum; LC: locus coeruleus; PB: parabrachial n.; FLM: Fasciculus longitudinalis medialis; LP: lateral pontine n. ; MP: medial pontine n.; SCv: ventral subcoeruleus ; TD: dorsal tegmental n.; DR: dorsal raphe.

Increases in noradrenergic tone are associated with motivation and an augmentation in behavioral drive. We therefore first tested the hypothesis that activation of noradrenergic system would increase song rate. Consistent with this hypothesis, stimulation of LC caused a significant increase in rate of undirected singing in all 4 of the implanted birds. LC stimulation caused a mean increase in singing of 2.30±0.65 songs/minute (figure 3b; hierarchical bootstrap t-test, p<0.0001).

**Figure 3.**
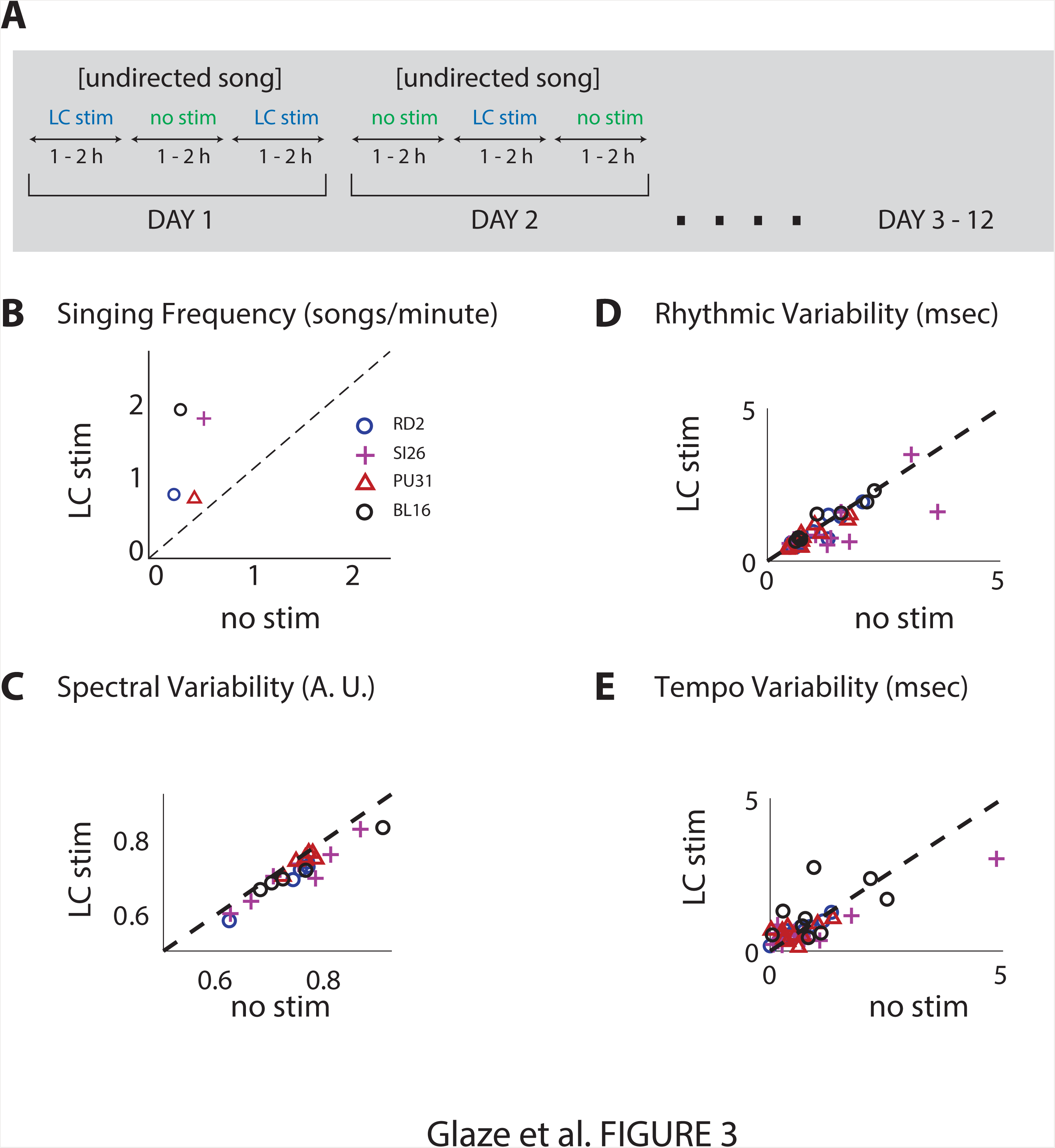
**Stimulation of the Locus Coeruleus increases singing rate and decreases the spectral and rhythmic variability of undirected song.** (A) Timeline of experimental paradigm (B) Scatterplot of the average number of songs produced per minute of recording from each of the 4 birds by experimental condition. Measures taken from the LC-stimulation condition fall along the vertical axis and those taken from the no-stimulation along the horizontal axis. (C-E) Scatterplots of spectral variability (C), rhythmic variability (D), and tempo variability (E) plotted for each song syllable (25 different syllables across the 4 birds), with axes as in (B). Syllables produced by each bird are indicated by different symbols or colors.

To determine the effect of LC stimulation on syllable variability, we used our novel “Euclidean variability” technique to quantify changes in spectral and temporal variability across all song syllables. Locus coeruleus stimulation caused profound changes in spectral variability, with stimulation during undirected singing decreasing spectral variability in all 4 birds by 4.25 ± 0.74% (figure 3c; hierarchical bootstrap t-test, p<0.0001).

Given the observed changes on various aspects of timing variability between directed and undirected song, we also were interested in addressing whether similar changes in timing variability would be observed during LC stimulation of undirected song. Indeed, compared to unstimulated song, LC stimulation caused a decrease in rhythmic variability across time intervals by 9.41±3.08%, with the direction of change persisting across all 4 birds (figure 3d; hierarchical bootstrap t-test, p<0.005). LC stimulation also indicated a trend of increased tempo variability by 95.57±35.7%, with the direction of change holding in 3 of 4 birds (figure 3e; hierarchical bootstrap t-test, p<0.05). Finally, LC stimulation also caused a modest but inconsistent increase in average tempo, with durations decreasing by 0.4±0.2% (p=0.178).

Thus, the directions of change effected by LC stimulation for singing rate and all of the spectral and temporal variability measurements were consistent with the directions of change observed by introducing a female (undirected to directed song) (Table 1).

**TABLE 1.** Summary of effects of NE manipulation on Song Features

|  | Relative to Undirected Song |  |  | Relative to Directed Song |
| --- | --- | --- | --- | --- |
|  | Direct | Undirect - LC Stim | Undirect - NE in RA | Direct - PHE in RA |
| Spectral variability | decrease (5.6%) | decrease (4.5%) | decrease (4.74%) | increase (8.2%) |
| Rhythmic variability | decrease (12%) | decrease (9.41%) | decrease (17%) | increase (12.3%) |
| Tempo variability | increase (56%) | increase (95%) | no change | no change |
| Tempo | no change | no change | no change | no change |
| Song rate | increase | increase | no change | no change |

<u>Legend</u>: LC stimulation of undirected song cause similar changes to song characteristics as the transition from undirected to directed song. In contrast, infusion of NE directly into RA only causes changes in spectral and rhythmic variability. The receptor antagonist PHE causes the exact opposite effect of NE when infused in RA while birds are singing directed song.

### Targeted increases of norepinephrine in RA decrease spectral and rhythmic variability of undirected song without increasing song rate

Because of widespread LC projections throughout the central nervous system, it remained unclear whether changes in song variability following LC stimulation were due to general changes in behavioral state or alternatively to a direct action on the vocal motor system. To test the hypothesis that norepinephrine has a direct effect on motor performance, we used a microdialysis probe to infuse norepinephrine (NE) directly into RA, a cortical motor nucleus that projects directly to brainstem vocal-respiratory centers and serves as the final common output of the forebrain song control circuit (figure 4a). Based on prior in vitro studies in brains slices (Solis and Perkel, 2006; Sizemore and Perkel, 2008), we hypothesized that infusion of NE would inhibit inputs from LMAN onto RA in a presynaptic fashion and thereby selectively reduce the acoustic variability imposed by the anterior forebrain pathway.

**Figure 4.**
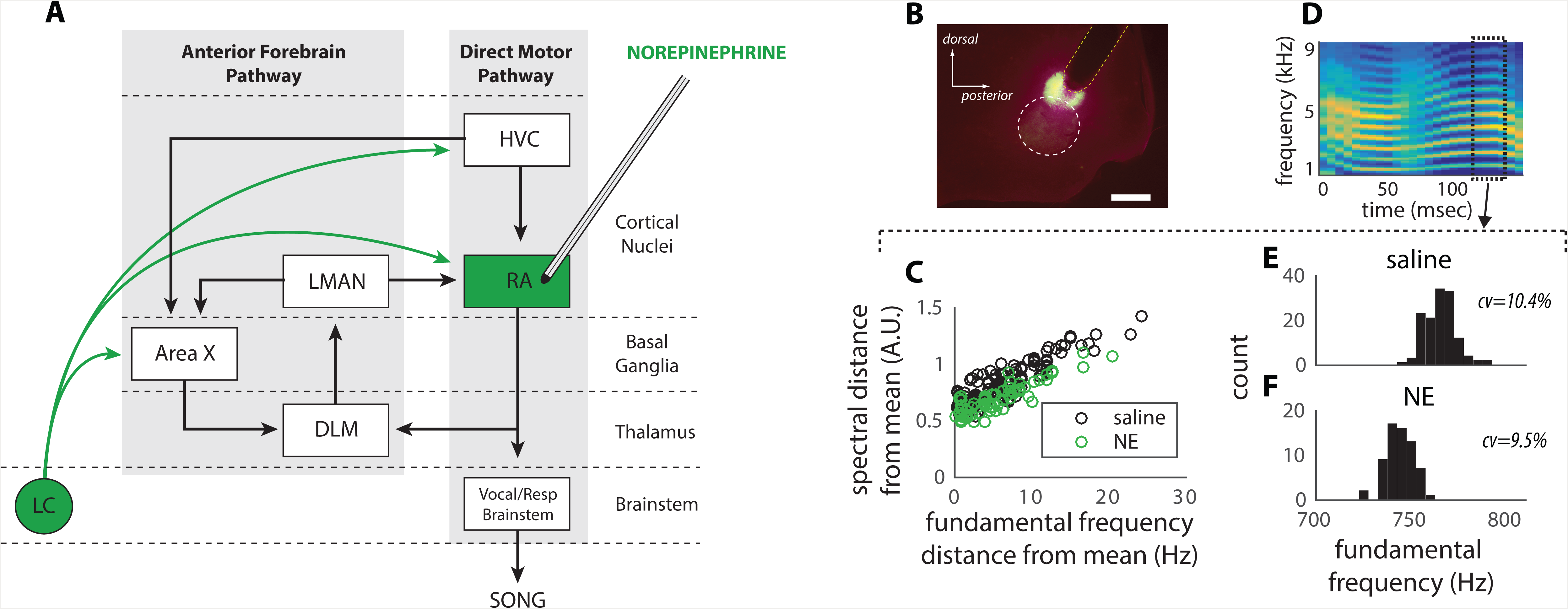
**Infusion of norepinephrine directly into nucleus RA.** (A) Illustration of the song system and probe implant into nucleus RA used for the microdialysis technique described in Methods. (B) Histology slide verifying anatomical location of probe implant in one bird by passing of fluorescently tagged muscimol through the probe after the experiment. Probes were typically placed in the dorsal pat of RA to prevent damage to the structure. RA contour is shown by the white dotted lined and probe by the yellow dotted line. Calibration bar = 500 µm. (C) Spectral variability scatterplot for a song syllable segment with a well-defined fundamental frequency as in Fig. 1C, with trial-by-trial spectral distance from the mean segment calculated using the novel method along the vertical axis and the more standard distance in fundamental frequency from the mean along the horizontal axis. Measurements during NE infusion are in green and those during saline are in black. (D) Spectrogram of the syllable analyzed in (C) along with key segment for which variability measurements were extracted. (E, F) Histograms of fundamental frequency during infusion of saline (E) and NE (F), with measured standard deviations indicated to the right. Overall, the analysis demonstrates a robust decrease in variability during confirmed NE infusion into RA that can be measured either with the novel or standard method.

A total of 8 birds were implanted with custom microdialysis probes targeted to the dorsal surface of RA. Care was taken not to place the probe directly into RA to avoid potential damage to RA (figure 4b). Birds implanted with microdialysis probes were infused on alternate days with either saline (control) or norepinephrine (NE) for periods ranging from 2 to 4 hours. Within each day, recording blocks were alternated between directed and undirected song. Because the current experiments were aimed at the hypothesis that elevations of NE in RA cause a decrease in song variability, songs were analyzed only under conditions of undirected song.

Similar to results obtained from LC stimulation, infusion of NE in RA caused a robust decrease in spectral variability compared to saline with a difference of 4.74±1.32 % and the direction of change holding in 7 of 8 birds (figure 4c-f, figure 5c; hierarchical bootstrap t-test, p<0.0001). NE infusion into RA also decreased rhythmic variability in 7 of 8 birds by an average 17.45±7.07% (figure 5d; hierarchical bootstrap t-test, p<0.01).

**Figure 5.**
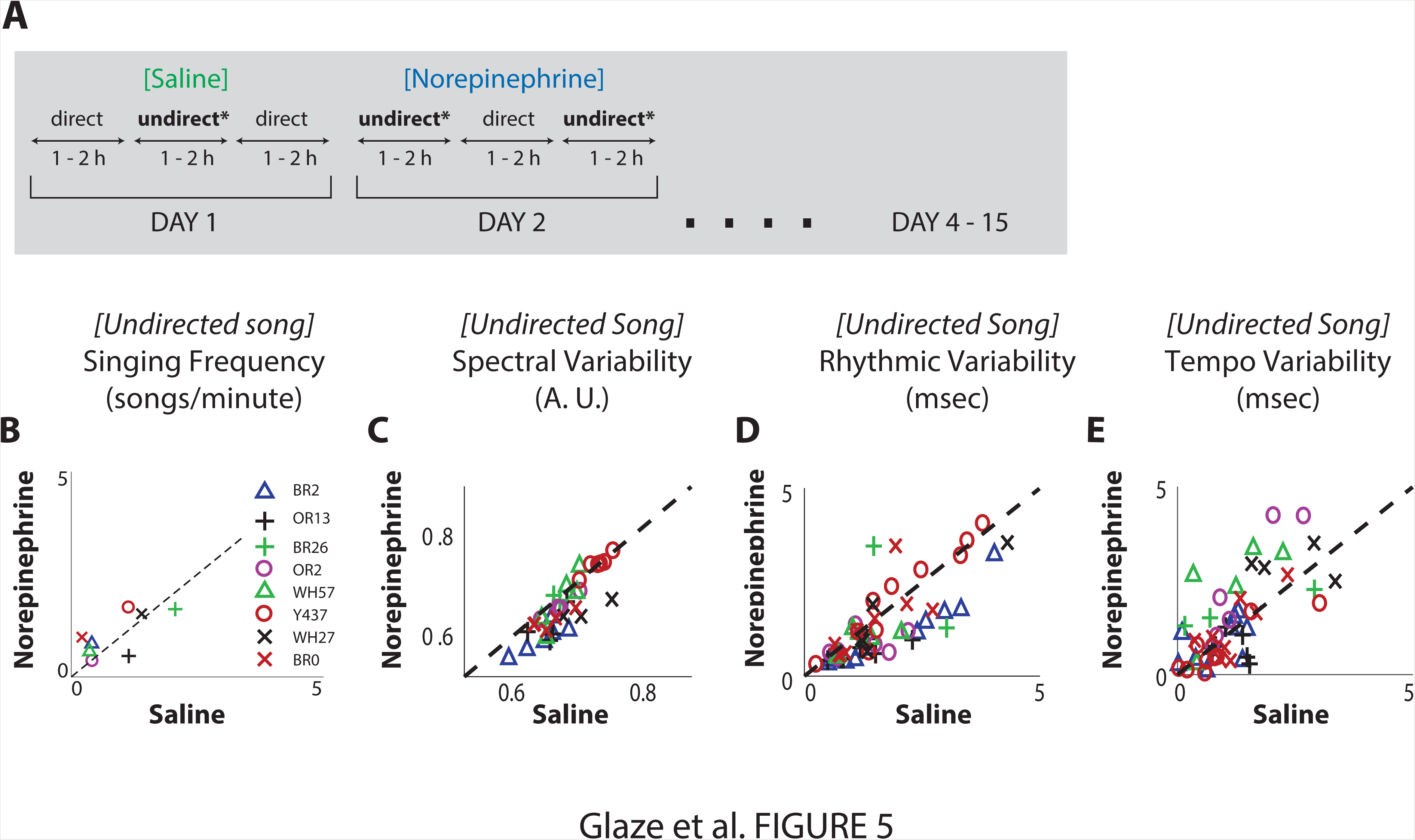
**Direct infusion of norepinephrine into nucleus RA decreases spectro-temporal variability of undirected song without changing song rate.** (A) Timeline of experimental paradigm. (B) Scatterplots of key song syllable parameters during infusion of NE (C, D, E) on the vertical axis and saline infusion condition on the x-axis. (B): Number of songs produced per minute of recording, with each symbol representing song rate for each of the 8 birds. (C) Spectral variability measurements using novel method; (D) rhythmic variability and (E) tempo variability. The analysis suggests that norepinephrine infusion into nucleus RA significantly decreased spectral and rhythmic variability in males singing undirected song.

Interestingly, in contrast to the effects observed following LC stimulation, elevation of NE in RA did not produce a significant change in tempo variability (figure 5e; increase of 37.66 ± 37.18%, with the direction of change holding in 5 of 8 birds, hierarchical bootstrap t-test, p=0.152) or in average tempo (1.12±0.5%; hierarchical bootstrap t-test, p=0.067). Similarly, norepinephrine infusion into RA did not affect song rate (0.308±0.262 more songs per minute compared with saline; figure 5a; hierarchical bootstrap t-test, p=0.121).

Taken together, these findings suggest that elevated levels of NE in RA only produce a subset of the effects observed following LC stimulation. While RA might be the target for several of the effects of norepinephrine (spectral and rhythmic variability), other target areas likely mediate effects such as song rate and tempo variability.

### Targeted infusion of RA with the NE receptor antagonist phentolamine increases spectral and rhythmic variability of directed song

Data obtained from LC stimulation and NE infusion into RA are consistent with a model in which the noradrenergic system gates the introduction of variability by the basal ganglia circuit into the motor system. We decided therefore to test the hypothesis that infusion of the non-selective alpha-adrenergic receptor antagonist phentolamine (PHE) during female-directed song, when NE should be elevated in RA, prevents NE-mediated suppression of LMAN input onto RA and thereby increases spectral variability.

We infused PHE directly into RA in four birds and recorded song both in the absence and presence of a female. Birds were infused with saline on alternate days (figure 6a). Because the current experiments were aimed at the hypothesis that elevations of PHE in RA might increase song variability, songs were analyzed only under conditions when males were singing directed song in the presence of a female. Song characteristics produced in the presence of PHE were compared directly to those produced during saline infusion. PHE infusion increased spectral variability of directed song in 3 of 4 birds, with an average increase of 8.22±3.19% (figure 6c; hierarchical bootstrap t-test, p<0.05). PHE infusion also increased rhythmic variability in 4 of 4 birds, with an average increase of 12.38±5.71% (figure 6d; hierarchical bootstrap t-test, p<0.05). Similar to the lack of effect of NE on tempo and tempo variability, PHE did not yield a consistent effects on tempo variability (figure 6e) or mean tempo although there was an insignificant trend toward decreasing tempo variability by 12.6±12.29% (trend in 3 of 4 birds, hierarchical bootstrap t-test, p=0.115) and increasing average tempo by 0.6±0.8 (direction of change in 2 of 2 birds, p=0.205). Also consistent with RA not playing a significant role in song rate, PHE infusion did not affect song production rate (1.065±1.241% in 2 of 4 birds; hierarchical bootstrap t-test, p=0.084; figure 6b).

**Figure 6.**
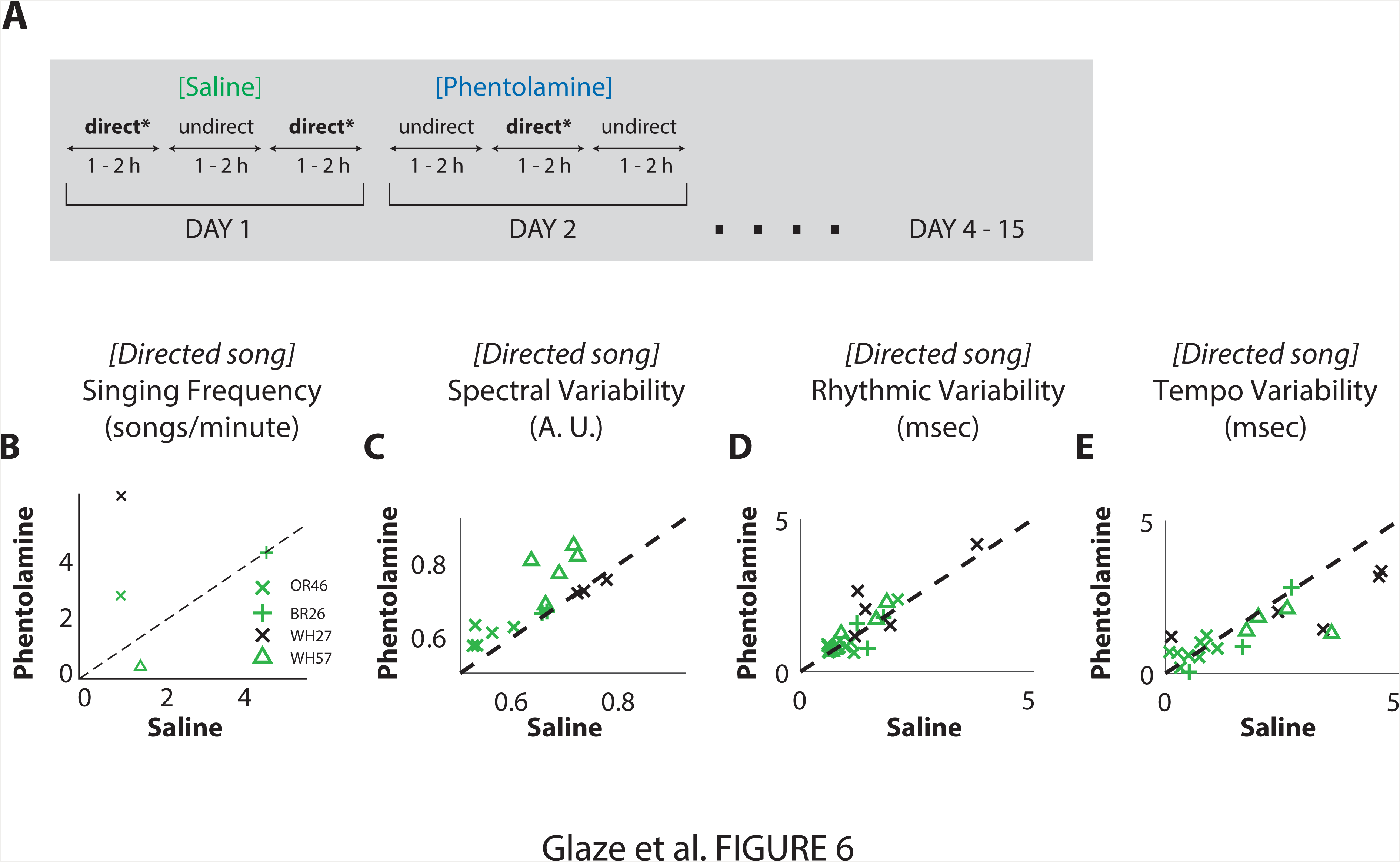
**Influence of direct infusion of phentolamine into nucleus RA increases spectral and rhythmic variability of directed song.** (A) Timeline of experimental paradigm. (B) Scatterplots of key song syllable parameters with infusion of the alpha-adrenergic receptor antagonist phentolamine (PHE)) on the vertical axis and saline infusion condition on the x-axis. (B) Number of songs produced per minute of recording, with each symbol representing song rate for each of the 4 birds. Three of these birds were also used for the NE infusion experiments (C) Spectral variability measurements using novel method; (D) rhythmic variability and (E) tempo variability. The analysis suggests that phentolamine infusion increases both spectral and rhythmic variability of female directed song.

Taken together, these findings suggest that preventing NE from blocking LMAN inputs onto RA by administration of the receptor antagonist PHE causes an increase in spectral and rhythmic variability of directed song.

## DISCUSSION

We investigated the influence of the noradrenergic system on the production of a learned and ethologically relevant vocal behavior in male zebra finches. Direct stimulation of the locus coeruleus in males singing alone (undirected song) recapitulated all measured features of female-present songs (directed song). These included (1) an increase in the frequency of produced songs, (2) greater spectral precision on fast timescales and (3) greater rhythmic precision on slower timescales. Interestingly, direct infusion of NE into RA, a motor nucleus in the song system that is analogous with primary motor cortex in mammals (Dugas-Ford et al., 2012), during undirected song resulted in greater spectral and rhythmic precision but did not increase song production or changes in tempo variability. Conversely, infusion of the non-selective adrenergic alpha-receptor antagonist (PHE) into RA during directed song singing resulted in decreased spectral and rhythmic precision without any statistically significant influence on song production frequency or tempo variability. The findings are summarized in Table 1.

The behavioral differences observed between the LC stimulation and direct, pharmacological manipulation in RA imply that song production rate and tempo variability is not mediated by noradrenergic modulation of RA and therefore likely is caused by direct or indirect effects of norepinephrine on other areas of the song control system. Possible targets for these additional effects on song might include the medial preoptic area, a midbrain nucleus that receives strong projections from LC, and which is known to project directly to the song system (Riters and Alger, 2004) and influence the motivation to sing (Riters and Ball, 1999; Riters et al., 2004). Our observation that a single brain structure (nucleus RA) is responsible for changes in rhythmic and spectral variability of song suggests that NE can have direct effects on the motor system that are distinct from a more generalized effect on arousal level and therefore provide support for the idea that LC projections contain a degree of anatomical specificity that allow them to differentially influence distinct cortical regions (Chandler et al., 2014a; Chandler, 2016).

### Implications for models of motor variability in the song system

The specificity of pharmacological effects suggests that increases in norepinephrine in RA are capable of generating a degree of behavioral precision that is similar to that observed during female-directed song. Previous work has shown that inactivation of LMAN, the output nucleus of the basal ganglia circuit onto RA, or blockade of its synaptic inputs, can cause a reduction in song spectral variability during the normally variable undirected song (Kao et al., 2005; Kao and Brainard, 2006; Stepanek and Doupe, 2010; Charlesworth et al., 2011). Because LMAN neurons show pronounced burst-like activity during undirected song (Kao et al., 2008), NMDA-dependent activation of RA neurons by these inputs is therefore thought to bias premotor activity within RA and cause the observed increase in variability of song output (Olveczky et al., 2005; Kao et al., 2008; Olveczky et al., 2011; Kojima et al., 2013). The present study therefore is consistent with a model, originally proposed by Sizemore and Perkel (2008), where increases in NE within RA suppress, in a presynaptic fashion, inputs from LMAN and thereby block the influence of the basal ganglia circuit and consequently decrease trial-to-trial spectral variability (Sizemore and Perkel, 2008).

The observation that activity in LMAN only shows variable bursting activity during undirected song (Kao et al., 2008), has generally led to the assumption that the more precise neural patterns produced during directed song do not play a significant role in generating variability in RA. It therefore is intriguing that infusion in RA of the NE receptor antagonist PHE caused a significant increase in song variability during female-directed song. One possible interpretation is that LMAN inputs during directed song carry more variability than previously appreciated and that this variability is normally suppressed by elevated levels of NE in RA during female directed song. Alternatively, it is possible that elevated levels of NE during directed song might influence song variability by acting directly within RA. Given that alpha-2 NE receptor manipulation can change RA neuron excitability of RA in vitro (Solis and Perkel, 2006), it is possible that addition of PHE to RA prevents the stabilizing effects of NE on the RA network in a way that results in increased spectral and rhythmic variability.

Although this study has identified RA as a major target for NE effects on song variability, it is likely that norepinephrine can also influence the generation of motor variability at other loci given that NE, perhaps in conjunction with dopamine (Leblois et al., 2010; Leblois and Perkel, 2012), acts directly on the basal ganglia (Area X) (Castelino and Ball, 2005) to reduce the generation of neural variability in pallidal neurons (Woolley et al., 2014).

### Implications for models of song timing and neural synchronization

Single-unit recordings in HVC show that RA-projecting neurons each produce a single burst of action potentials that is precisely time-locked to distinct epochs in the song (Hahnloser et al., 2002; Lynch et al., 2016) with correlational data suggesting that these bursts control the timing in RA song premotor output (Fee et al., 2004). Because cooling of HVC, but not RA, slows overall song tempo (Long and Fee, 2008) it has been suggested that HVC controls much of song timing (Fiete et al., 2007; Long and Fee, 2008; Long et al., 2010). The observed effects of norepinephrine manipulation in RA on rhythmic variability of song are inconsistent with the simplest interpretation of these models and suggest that song timing control might be implemented in a more distributed manner across the song system (Ashmore et al., 2005; Hamaguchi et al., 2016; Schmidt and Goller, 2016): while HVC may influence RA timing, RA timing may also influence the timing of spikes in HVC and elsewhere. Such an influence could be direct, for example mediated by unilateral projections to HVC from the dorsal region of RA (dRA) (Roberts et al. 2008) or indirect, as through recurrent bilateral inputs from the RA-PAm-UVa-HVC pathway (Ashmore et al., 2005; Hamaguchi et al., 2016). Interestingly, our findings leave open an interesting question as to why cooling RA does not have any apparent influence on song given that NE and PHE infusion cause changes in song timing. Perhaps cooling has a differential influence on the gain of bursting activity in neurons in the dRA that preempts influence on spike timing in HVC. In contrast, pharmacological manipulations of NE within RA might keep the gain relatively intact while perturbing the statistical burst properties of RA neurons, which could in turn propagate through HVC and the rest of the song system to modify song timing.

### Implications for the role of noradrenergic system in other systems

The noradrenergic system has been proposed to facilitate behavioral optimization (Aston-Jones and Cohen, 2005a, b) as well as explore-exploit motor learning. Under this theory, tonic increases in LC spiking and the associated increases in norepinephrine enhance “distractibility” and promote exploration over a wide set of possible salient stimuli or behaviors. In contrast, phasic LC spiking, and as a consequence the more moderate increase in released NE, promote more focused task engagement and exploitation of actions previously learned as optimal for reward maximization. To date, this theory has been tested primarily in a discrete action space where optimal behavior is defined as a decision among a discrete set of options, such as in foraging tasks in which the subject must choose, within a given trial, among one of several options with uncertain reward value (Kalwani et al., 2014; Bouret and Richmond, 2015) . In the zebra finch, we tested the role of NE in song production, which provides an interesting example of behavioral optimization. Here the optimal behavior is not the choice of a specific action but rather the choice of temporal and spectral motor commands in a sequenced, hierarchically organized composition of actions (Glaze and Troyer, 2006) that are produced in a continuous feature space (Tchernichovski et al., 2001). It is thus unclear what the analogue to phasic spiking would be in this context. In the current study, we manipulated NE levels either by direct infusion in RA or by imposing tonic 1Hz stimulation in LC and therefore could not direct address phasic vs. tonic changes given that our manipulations caused sustained increases in NE levels. In future experiments it will be important to manipulate NE levels in a more phasic fashion as well as record from single units from LC during song production to gain greater insight into how the LC network promotes exploitation of learned behavioral sequences. More generally, the influence of NE on signal-to-noise ratios in individual neurons shown in songbirds (Solis and Perkel, 2006; Sizemore and Perkel, 2008) and mammals (Berridge and Waterhouse, 2003) suggest that while the noradrenergic system appears to have a very diverse set of influences across the central nervous system, the common principle may simply change the relative gain of influence of neural signals linked with the greatest salience or utility (Aston-Jones and Cohen, 2005b; Mather et al., 2015).

## MATERIALS AND METHODS

### Subjects

Male and female zebra finches (*Taeniopygia guttata*) ranging from 120 to 250 days were used in this study. Animals were housed in same sex cages until the experiment commenced. Test males with either stimulating electrode or dialysis probe implants were housed alone in sound proof chambers. All the birds were housed on 14h Light: 10h Dark and given food and water *ad libitum*.

### LC stimulation

All implantation surgeries were performed on birds anesthetized with a mixture of ketamine (40mg/kg; Phoenix Pharmaceuticals, Belmont, CA) and xylazine (8 mg/kg; Phoenix Pharmaceuticals, Belmont, CA). A total of 17 male zebra finches were implanted with stereotrodes (25 *μ*m diameter nichrome wire). Although full length experiments were performed on most of the birds, several birds either died or lost their heads caps before we could perform electrolytic lesions to confirm electrode location. Of the remaining birds where we placed electrolytic lesions (60 *μ*A for 10 seconds), we were able to confirm that electrodes were placed in both the left and right LC in 4 birds (Fig 2b). Electrode location was assessed visually in fresh unstained tissue with the criterion that the center of the lesion site be within the anatomical confines of the LC. Implant placement was guided by a combination of stereotaxic coordinates and electrophysiological landmarks. Implanted birds were tethered to a mercury commutator that was connected to an A-M systems stimulator (Model 1800). Because the goal of these experiments was to test the effect of LC stimulation on undirected song, recordings were always made from male birds placed in a cage without a female. A typical experimental day consisted of LC stimulation periods ranging from 1 to 2 hours that were interleaved with periods of non-stimulation of roughly the same duration for approximately 6 hours each day. Experiments were designed so that the order of stimulation and non-stimulation periods was reversed on each experimental day (Figure 3a). Stimulation parameters consisted of tonic 1 Hz, 150 μA bilateral, biphasic pulses of 400 μsec duration. Such stimulation rates are consistent with median firing rates observed in LC of awake behaving rodents (Fazlali et al., 2016). Each of the implanted birds was subjected to this pattern of LC stimulation for 3 to 12 days.

### Reverse microdialysis

A total of nine birds received microdialysis probe implants in each RA that were placed based on a combination of electrophysiological characteristics and stereotaxic coordinates. Probes were aimed to the dorsal edge of RA to minimize placing the cannula directly into RA and damaging the structure (figure 4b). Unlike injections, dialysis probes allow drugs to diffuse out of the cannula without adding any volume to the targeted area. Probes were manufactured according to (Olveczky et al., 2011) using Spectra/Por^®^ in vivo microdialysis hollow fibers (Spectrum Labs, Ranchos Dominguez, CA) attached to polyimide tubing to make probes that had a final overall diameter of approximately 250 µm. The dialysis membrane had a 13 kD cutoff which was sufficient to allow for diffusion of NE and phentolamine (PHE) into RA. Probes placed in the left and right RA were connected to each other in series and drugs were infused through FEP tubing (CMA/Microdialysis, Holliston, MA) that was connected to the probes using a syringe pump (WPI, Sarasota, FL). Drugs and saline were infused through the probes at 2 to 4 ml/hour. Probe placement in RA was verified by passing Fast Green dye or fluorescently tagged muscimol (BODIPY TMR-X muscimol conjugate, ThermoFisher Scientific) through the probe at the end of each experiment for approximately 30 minutes. Dye spread around the probe tip was combined with histological processing to verify the cannula tip was placed on the dorsal edge of RA. Norepinephrine (NE; Sigma-Aldrich) and the nonselective alpha-adrenergic receptor antagonist phentolamine (PHE; Sigma-Aldrich) were dissolved in physiological saline to a final concentration of 15 mM for NE and PHE. Each testing day was divided up into 2 hours of singing in one context (e.g. female present) followed by 2 hours of singing in the other context (e.g. female absent). We recorded between 2 to 4 sessions (of 2 hours each) per day, and alternated the order of social context (i.e. the presence or absence of a female) across consecutive days. Only one condition (drug or saline) was used on any given day, and this was alternated across consecutive days (Figures 5a and 6a). Overall duration of the experiments lasted 4 to 15 days.

### Song recording and selection

Songs were digitized at 44.1 kHz and collected as wav files using Song Analysis Pro (SAP) (Tchernichovski et al., 2000). Files were automatically screened in SAP for those most likely to contain song. The resulting wav file data were then analyzed using custom software written in Matlab (Mathworks, Natick, MA). As described earlier, experimental conditions consisted of males housed either in isolation or in the presence of a female. We defined undirected songs as those songs produced when the male was singing by himself in the absence of a female. Consistent with other studies (Woolley and Doupe, 2008), we defined directed songs as those songs produced by the male during the first 10 minutes after he was exposed to a female. Songs produced after the first 10 minutes often contain a mixture of directed and undirected song and therefore were not further analyzed in our studies because of the ambiguity of defining the songs as being directed or undirected.

#### Song Analysis

The songs of each bird in each experimental condition were analyzed by (1) using a fully automated custom-written user-trained pattern matching algorithm that identifies song syllables across the entire data set and quantifies the most common sequence of identified song syllables, (2) quantifying song characteristics such as singing rates and the spectral and tempo variability of each syllable for each bird in each experimental condition, and (3) computing statistics using a bootstrap procedure. Each of these steps are described in more detail below:

##### 1. Song syllable and motif identification

Each wav file was divided into sound clips using threshold crossings in the log-power of the signal. We then selected a random sample of 500-1000 clips from the resulting set (across all wav files). Individual sound clips were transformed into spectrograms with a 512-point window slid forward in 256-point steps and filtered between 500 and 9,000 Hz. Using a custom written GUI in Matlab, each clip spectrogram was then manually classified as a particular song syllable (e.g. syllable “a”, syllable “b”, etc.). Manual classification was based on a combination of visual inspection, syllable playback, and statistics computed in the GUI that includes average fundamental frequency, Wiener entropy and Euclidean distance to other syllables in the sample.

The resulting “training set” was then used to automatically identify instances of each song syllable across the entire wav file set specific to each bird and experimental condition. Spectrograms for each syllable were reduced to 5 dimensions along the frequency axis using Principal Components Analysis (PCA) to construct a basis vector for each syllable. This information was then used to concatenate vectors for sound clip spectrograms into a single matrix and associated with the 5 largest eigenvalues of the covariance matrix. Each clip was then projected onto the new basis to yield a 5 x N matrix, where N is the number of time points in the original clip spectrogram. Each wav file across the entire data set was then treated with the same sequence of transformations as the sound clips, i.e. filtered between 500 and 9000 Hz, transformed into spectrograms and reduced to 5 dimensions along the frequency axis using the same principal components extracted from the training set. A sliding cross-correlation was then computed between each reduced clip and the reduced wav file, and a custom “sliding” k-nearest neighbors algorithm (Bishop, 2006) was used to identify instances of each syllable class. Specifically, for each time-point in the wav file, the 11 highest cross-correlation scores were collected along with associated clip classes; a class (or “match”) was then attached to a given time-point if at least half of the highest scores were associated with that class. In cases where two of the matches overlapped, the one with the highest average score was chosen.

After syllables were classified, the most common song motif for each bird was identified using an automated algorithm that compiled all unique syllable sequences and picked the sequence with at least 2 syllables that was most common. Syllables were considered adjacent in the same sequence if the silent gap between them was <200 msec of average duration. The resulting data set consisted of an average of 200 motif renditions per bird per condition (range 59-484) and a median 5 syllables per motif (range 3-9).

##### 2. Quantifying song characteristics

###### Singing rate analysis

To probe the frequency of singing per unit time we used an automated algorithm that searched for at least two classified song syllables in sequence (i.e. separated by <200 msec in average silent gap duration) and calculated the frequency of occurrence per total time of song recording for that bird.

###### Syllable spectral analysis

Each unique syllable’s rendition-to-rendition variability was analyzed within the most common motif sequence for that bird. <u>First</u>, each syllable’s rendition was normalized by the total amount of power in the acoustic signal and mapped to a time-base common to all renditions of that syllable using dynamic time-warping (Rabiner et al., 1989). Specifically, syllable spectrograms were aligned and averaged to yield a single template to which all renditions were mapped. <u>Second</u>, the resulting mapped spectrograms were averaged and the process was repeated, resulting in a “mean” syllable that had greater fidelity than the original. <u>Third</u>, we then calculated the “spectro-temporal deviation” of each syllable rendition as the Euclidean distance between rendition *j* of syllable *k* and the mean of syllable *k* as:

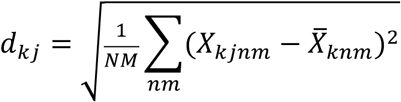

Where *N* and *M* are the number of time and frequency bins, *n* and *m* index time and frequency, *X* is the spectrogram of a syllable rendition, and 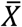 is the averaged syllable template described above. <u>Fourth</u>, we then computed total syllable-specific “spectro-temporal variability” (within a given experimental condition) as:

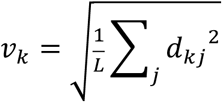

Where *L* is the total number of renditions for that syllable.

The values of *d_kj_* and *v_k_* can be thought of simply as broad measures of how “different” a syllable sounds from its average rendition, encompassing variability in the fundamental frequency of a vocalization with well-defined pitch, as well as variability in more complex sound features such as those found in amplitude-modulated notes and sweeps.

Previous investigations into song learning and contextual influences on variability have focused on the fundamental frequency (FF) of song elements that have a well-defined pitch (e.g. “harmonic stacks”). The more general measure of spectral variability we use here makes fewer assumptions about the nature of the signal. In a subset of syllables with a well-defined fundamental frequency we verified that the measure, which is represented in arbitrary units (A.U.), was correlated with estimates of FF (e.g. figure 1c,d).

###### Timing variability analysis

To measure precise differences in timing variability we re-calculated song spectrograms using 256-point windows slid forward in 128-point steps and calculated for each song syllable a template of the time-derivative using methods described previously (Glaze and Troyer, 2007, 2013). We then applied a dynamic time-warping algorithm (Rabiner et al., 1989; Glaze and Troyer, 2006, 2007) to measure syllable onset and offset times, yielding a sequence of syllable onset and offset times for each bird and associated motif, defining “time intervals” as the duration from either syllable onset to offset (i.e. syllable duration) or syllable offset to the onset of the next (silent gap durations). Finally, we decomposed rendition-to-rendition timing variability across the sequence into variations linked with global tempo modulations and variations independent of both tempo of measurement error in onset and offset times using a generative model described previously (Glaze and Troyer, 2012) (figure 1e,g). Specifically, we wrote time interval *i* in rendition *n* as:

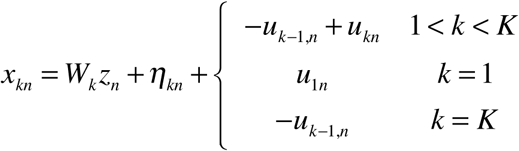

Where *x* is the time interval, *z* is the (0-mean, unit-variance) tempo for that rendition, *W* is the relative contribution of interval *x* to this shared tempo, *u* is jitter in the onset/offset time measurement that does not contribute to overall motif duration, *K* is the total number of time intervals in the motif, and *η* is the timing variable independent of both tempo and jitter, influencing overall song duration independently of the other intervals. This last component may be thought of intuitively as “rhythmic variability” because it significantly changes the proportional durations of each sequence element significantly more than the global component and cannot in principal be attributed to measurement error because it influences the timing of every subsequent element (Glaze, 2008; Glaze and Troyer, 2012).

Song sample sizes for this analysis were significantly smaller compared with previous studies estimating these components, making estimation especially susceptible to sample error using maximum likelihood techniques. We thus used a modified version of an expectation-maximization (EM) algorithm previously developed to make component estimates (Glaze and Troyer, 2012). The central difference here was in the “maximization step”. Here estimates at each iteration of the algorithm were based on maximizing the posterior probability of each parameter given the data and (1) a 0-mean, unit-variance Gaussian prior probability distribution constraining estimates of W and (2) an exponential prior probability distribution constraining estimates of the total variance in *η* (i.e. rhythmic variability). Component estimates corroborated previous findings from the maximum-likelihood based method that global timing changes were indeed dominated by a shared positive correlation in trial-to-trial variability among all time intervals, with ~94% of tempo weights (*W*) being positive, and roughly the same amounts of variability in the tempo and rhythmic variability components (medians of 0.869 and 1.292 msec respectively).

##### 3. Statistical analysis

Analysis was focused in 5 key measurements described above: (1) rate of singing, (2) rendition-to-rendition spectral variability in each syllable, (3) average duration of each syllable and silent gap, (4) magnitude of tempo variability contributed by each syllable and gap, and (5) rhythmic variability contributed by each syllable and gap independently of the rest. Analysis was performed for each of these measures across different experimental conditions: (i) presence of a female, (ii) stimulation of LC (iii) infusion of NE into RA or (iv) infusion of PHE into RA.

We tested for the significance of each of these effects using a bootstrapping procedure (Davison and Hinkley, 1997) that accounted for random sampling of both individual birds and syllables rather than treating each syllable independently. Specifically, for each hypothesis test we constructed 1000 bootstrapped samples of the mean difference in measurement across conditions by re-sampling individual birds and song renditions with replacement. A single bootstrapped mean was thus calculated as the average across a set of differences obtained by (1) randomly sampling a bird, (2) randomly sampling motif renditions for that bird, (3) computing song rate, syllable variability and timing variability over that sample, (4) repeating steps (1) and (3) until the total number of birds for the given experiment (e.g. LC stimulation) had been selected, and (5) calculating the mean of paired differences between conditions. We then calculated standard errors on mean differences as the standard deviation of the resulting distribution of differences. We then defined statistical significance with the percentage of all bootstrapped samples falling below/above 0 per each hypothesis, using a two-tailed 5% threshold (i.e. significant difference if 97.5% of samples either fall above or below 0). In our calculations of song rate, bootstrapped samples of differences in rate of song production were calculated by taking the total number of vocal sequences produced across the random bird sample and dividing by the total amount of song recording time in order to avoid the disproportionate influence of birds with relative small song recording time durations. Throughout, we refer to this as a “hierarchical bootstrap test” and report p-values based on the percentage of bootstrapped samples falling below/above 0 along with the mean and standard deviation of the sample distribution (the latter yielding standard errors on the differences). We report the consistency of effect differences across birds using median parameter differences by bird where parameters were calculated across all samples for each bird.

## Acknowledgements

We thank David Perkel and members of the Schmidt lab for reading an earlier version of this manuscript. We also thank the three reviewers for insightful comments that helped significantly improve the manuscript. We are also indebted to Richard Hahnloser and Janie Ondracek for teaching us how to make the microdialysis probes.

